# Hoxa10 mediates positional memory to govern stem cell function in adult skeletal muscle

**DOI:** 10.1101/2020.07.16.207654

**Authors:** Kiyoshi Yoshioka, Hiroshi Nagahisa, Fumihito Miura, Hiromitsu Araki, Yasutomi Kamei, Yasuo Kitajima, Daiki Seko, Jumpei Nogami, Yoshifumi Tsuchiya, Narihiro Okazaki, Akihiko Yonekura, Seigo Ohba, Yoshinori Sumita, Ko Chiba, Kosei Ito, Izumi Asahina, Yoshihiro Ogawa, Takashi Ito, Yasuyuki Ohkawa, Yusuke Ono

## Abstract

Skeletal muscle stem cells (satellite cells) are distributed throughout the body with heterogeneous properties that corresponds to region-specific pathophysiology. However, topographical genes that have functions remain unidentified in satellite cells of adult muscle. Here, we showed that expression of Homeobox (Hox)-A cluster genes, key regulators of the embryonic body plan, was robustly maintained in both muscles and satellite cells in adult mice and humans, which recapitulates their embryonic origin. We observed that regionally specific expressed Hox genes were linked to hypermethylation of the Hox-A locus. We examined *Hoxa10* inactivation in satellite cells and found it led to genomic instability and mitotic catastrophe, which resulted in a decline in the regionally specific regenerative ability of muscles in adult mice. Thus, our results showed that Hox gene expression profiles instill the embryonic history in satellite cells as positional memory, potentially modulating the region-specificity in adult skeletal muscles.

## INTRODUCTION

Skeletal muscle is the contractile tissue that occupies approximately 40% of an individual’s body weight and is distributed throughout the body. Skeletal muscle has functionally heterogeneous properties that are specific to muscle type. Limb and trunk muscles function in body posture, locomotion, and respiration; while craniofacial muscles mainly control facial expression, speech, feeding activity, and eye movement. Most skeletal muscles of the trunk and limbs are derived from myogenic precursor cells that migrate from somites during embryonic development in vertebrates, whereas head musculature including masseter (MAS) muscles arises mainly from the cranial mesoderm *(1)*.

Resident muscle stem cells, also called satellite cells, are located between the basal lamina and the plasmalemma of myofibers *(2–4)*. Satellite cells provide myonuclei for postnatal muscle growth and hypertrophy, repair, and regeneration in adult muscle *(5)*. Recent studies revealed that satellite cells are a functional heterogeneous population in different muscles, and dependent not only on fiber-types but also on embryonic origin *(3, 4)*. We previously reported that satellite cells isolated from extensor digitorum longus (EDL) muscle, which originate from somite, differ in their fate-determination dynamics during proliferation, differentiation, and self-renewal compared with those from MAS muscle that originate from the cranial mesoderm *(6)*. Satellite cells in the nasopharynx muscle of the head constitutively proliferate and contribute to myonuclear turnover without muscle injury *(7)*, whereas these phenomena are rarely observed in limb muscles. The heterogeneity in the satellite cell population among muscles may therefore be related to region-specific pathophysiological phenotypes of muscle diseases *(8)*.

There are distinct genetic networks in pre-myogenic progenitors between head and trunk/limb muscles during embryonic development *(1)*. Paired-box protein Pax3 regulates limb myogenesis, whereas double-ablation of T-box transcription factor Tbx1, bicoid-related homeodomain transcription factor Pitx2 and musculin/Tcf21 play important roles in the specification of head muscle progenitor cells *(9–14)*. Satellite cells also share the ontogeny of the muscle, and topographic genes expressed in satellite cells in the head are distinct from limb *(6, 9, 15)*. Although satellite cell heterogeneity among muscles may link to the body-region-specific pathological phenotypes in muscle diseases *(3, 4, 8)*, topographical genes that have functions remain unidentified. In addition, the epigenetic regulation of topographical gene expression in muscles and satellite cells is largely unknown. DNA methylation is a stable epigenetic mark that associates with silencing and upregulation of gene expression *(16)*.

Here, we performed the global DNA methylome and transcriptome analysis on different muscles and their associated satellite cells from adult mice to comprehensively uncover region-specific epigenetic and transcriptional properties. Hox genes are well known as the master regulators of the animal body-plan that guide morphogenesis during development by controlling the genetic network program for temporal and spatial development of tissues and organs *(17)*. Vertebrates have 13 Hox paralogs with four gene clusters (A-D). We found that adult muscles and their satellite cells expressed Homeobox (Hox)-A cluster genes whose expression patterns reflected the anatomical location and embryonic history. We also identified the *Hoxa10* gene as a topographically functional gene: postnatal deletion of *Hoxa10* in satellite cells led to genomic instability and abnormal mitosis, resulting in a decline in the regionally specific regenerative ability of muscles. Thus, our findings suggest that Hox gene expression profiles are maintained in regionally specific patterns as functional positional memory in both adult muscles and their satellite cells.

## RESULTS

### Hox-A cluster locus is hypermethylated in limb muscle and associated satellite cells

To compare regionally specific DNA methylation between the head and limb muscles, we conducted the DNA methylome analysis using MAS muscle tissue from the head and tibialis anterior (TA) muscle tissue from the hindlimb in adult mice (Figure 1A). We performed the Microarray-based Integrated Analysis of Methylation by Isoschizomers (MIAMI)*(18)* and found that seven probes for Hox-A cluster genes including *Hoxa2, Hoxa3,* and *Hoxa5* were listed as the top ten probes of DNA hypermethylated genes in TA muscle compared with MAS muscle (Figure 1B). To validate the reliability of our MIAMI data, we performed additional DNA methylome analysis using the post-bisulfite adaptor tagging (PBAT)-mediated targeted methylome sequencing *(19, 20)* on samples from MAS and TA muscles and their associated satellite cells (Figure 1A). DNA methylation levels of all the CpG sites (Figure 1C), PCA analysis (Figure 1D), and unsupervised hierarchical clustering (Figure 1E) indicated similar methylation patterns among the samples (n= 3 mice). We performed gene ontology (GO) enrichment analysis based on hyper- or hypo-DNA methylated genes, which were described with embryonic development-related GO terms, such as anterior/posterior pattern specification, embryonic skeletal system morphogenesis, proximal/distal pattern formation, and embryonic hindlimb morphogenesis (Figure S1A-S1B). Vertebrates have 13 Hox paralogs with four gene clusters (A-D), which are located on different chromosomes (Figure 1F). Corresponding to the MIAMI data, only the Hox-A cluster was remarkably hypermethylated in both muscle tissues and satellite cells from TA muscles compared with satellite cells from MAS muscles (Figure 1G). DNA methylation profiles of the Hox-A locus demonstrated that Hox-A genes, including *Hoxa2, Hoxa3,* and *Hoxa5,* are hypermethylated in TA muscle and its associated satellite cells (Figure 1H).

**Figure 1.**
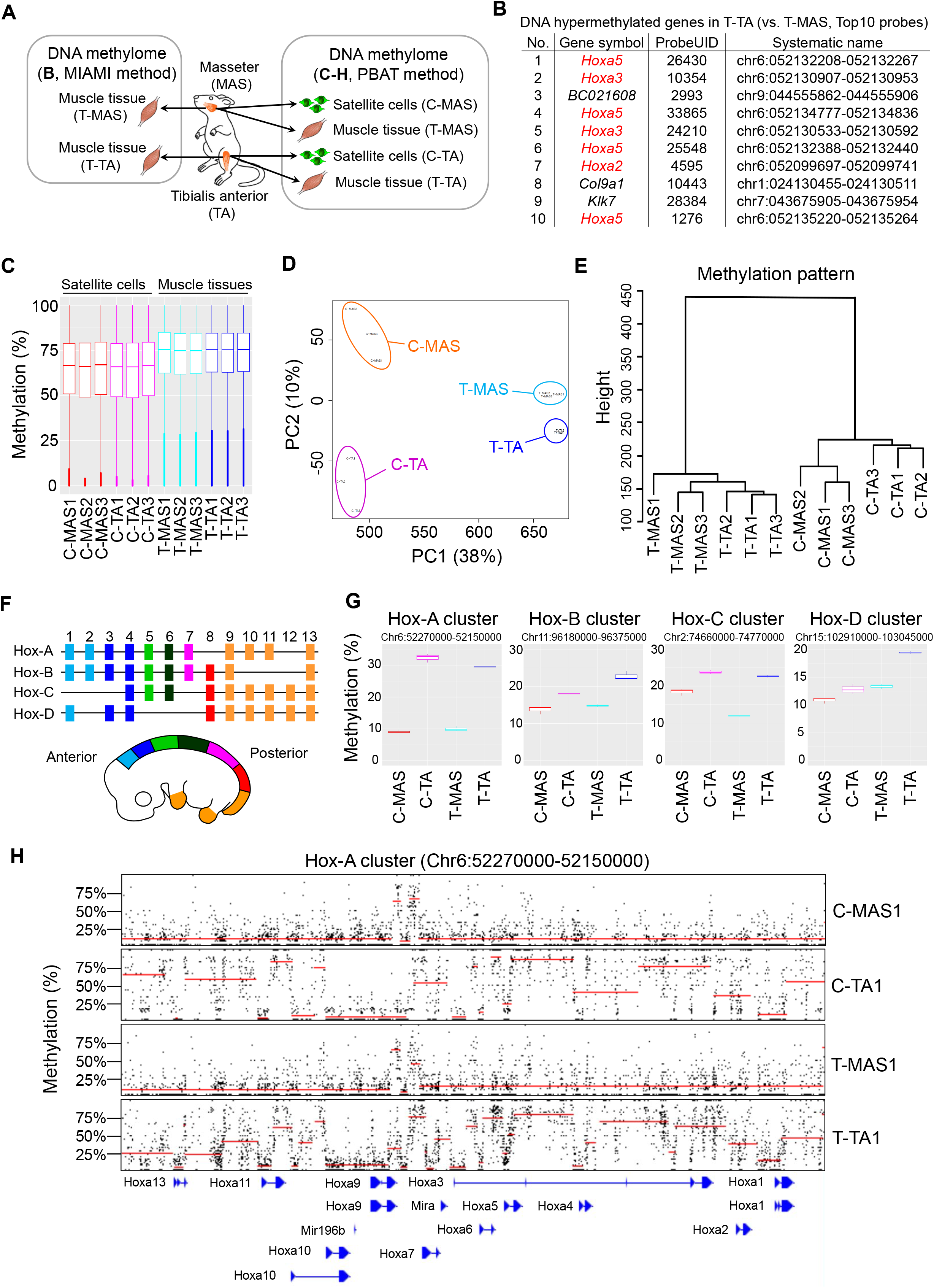
DNA methylation in head and limb muscles and associated satellite cells. (A-H) Schematic illustration of DNA methylome analysis in adult mice. Global DNA methylation status in MAS or TA muscle tissues and their associated satellite cells was evaluated using the MIAMI method (B) and PBAT method (C-H). (B) The top ten probes of DNA hypermethylated genes identified from TA muscle (vs. MAS muscle). (C) Boxplot of mean CpG methylation level of 1-kb sidling windows. (D) PCA analysis for DNA methylation. (E) Hierarchical clustering using Euclidean distance across the entire sample set. (F) Schematic illustration of Hox cluster genes in the mouse embryo. (G) DNA methylation profiles for each Hox cluster. (H) DNA methylation profiles for Hox-A cluster. The black dots indicate the methylation levels of individual cytosines in CpG context covered by at least 5 reads. The red lines indicate the domains calculated with a changepoint detection method *(40)* and their mean methylation levels are reflected in their positions in y-axis. (B, n= 1 mouse; C-H, n= 3 mice).

### Hox gene expression is maintained in adult muscles and their associated satellite cells

Hox-A and Hox-C cluster genes are known to be expressed in satellite cells in adult mice *(21–23).* Given that the hypermethylation status at Hox-A cluster locus was observed in limb muscles (Figure 1), we next determined whether the Hox gene expression profile differed in various muscles and their associated satellite cells. We performed a microarray using different muscle tissues in adult mice, and found a unique expression pattern of Hox-A and Hox-C cluster genes (Figure 2A). A recent study reported that DNA hypermethylation on the HOX-A locus is positively correlated with upregulation of HOX-A cluster genes in human glioblastoma (51), suggesting that DNA hypermethylation links upregulation of HOX-A cluster genes. Corresponding to this finding, we showed that Hox-A and Hox-C cluster genes were abundantly expressed in both fore- and hind-limb muscles, which originated from somite in development. However, only a few genes in Hox-A and Hox-C clusters were detected in MAS, digastric (DIG), and trapezius (TRP) muscles that originated from the cranial mesoderm *(6, 9, 15, 24)* (Figure 2A). We used four representative adult muscle tissues and their associated satellite cells, which included MAS and TRP with cranial mesoderm origin, supraspinatus (SS) and TA with somite origin, TRP and SS that are anatomically adjacent but distinct origins (Figure 2B–2C). Genes that modulate the limb skeleton pattern are known to be regulated by the Hox paralogs *Hox9–13 (17)*. Quantitative PCR (qPCR) analysis revealed that several Hox-A cluster genes such as Hoxa9–13 were detected in SS and TA limb muscle tissues, and appear to be expressed in a proximal-distal dependent manner (Figure 2B). TRP muscle abundantly expressed *Hoxa5–7* but not *Hoxa9–13* genes, whereas Hoxa5–13 genes were undetected in MAS muscle (Figure 2B). Muscle tissues and their associated satellite cells similarly expressed Hox-A cluster genes (Figures 2B and 2C). Single cell RNA-seq analysis further revealed that Hox-A and Hox-C cluster genes were detected in EDL hindlimb-derived satellite cells in culture (Figure 2D).

**Figure 2.**
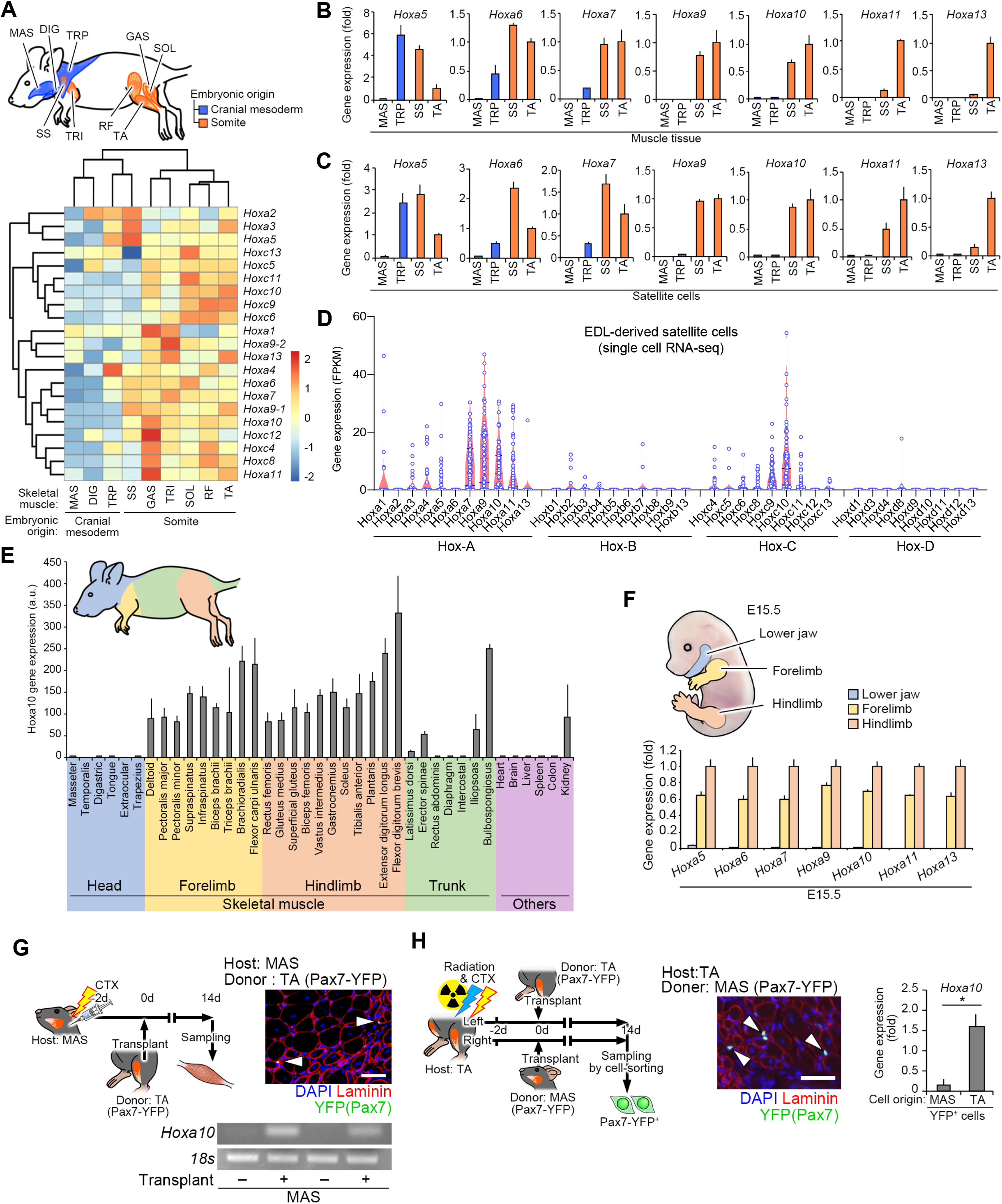
Hox gene expression pattern in different muscles and associated satellite cells. (A) Schematic illustration of cranial mesoderm-derived head muscles (blue) and somite-derived limb muscles (orange) in adult mice. Heatmap generated from microarray analysis demonstrated expression patterns of Hox-A and Hox-C cluster genes. (B and C) qPCR analysis of *Hoxa5–13* genes expressed in muscle tissues (B) and cultured satellite cells (C) in adult mice (n= 4 mice). Data represent means ± SEM. (D) Single cell RNA-seq analysis of Hox gene expression in cultured satellite cells isolated from EDL limb muscles from adult mice (n = 74 cells). (E) qPCR analysis of *Hoxa10* gene expression in muscle tissues, including head (blue, forelimb (yellow), hindlimb (orange), and trunk (green), and other organs (purple) from adult mice (n= more than 3 mice for each). Data represent means ± SEM. (F) qPCR analysis of *Hoxa5–13* gene expression in the lower jaw (blue), forelimb (yellow), and hindlimb (orange) at E15.5 (n= 5 mouse embryos). (G) Schematic illustration of the transplantation assay. TA muscle homogenates (Donor: *Pax7-YFP* mice) were engrafted into MAS muscle (Host: WT mice) pre-injured with a CTX injection. Representative immunohistochemical image of MAS transverse sections for laminin (red), YFP (Pax7, green) and DAPI (blue). Scale bar, 50 μm. A representative image of an agarose gel electrophoresis containing PCR products from *Hoxa10* and *18S* genes 14 d following transplantation (n= 5 mice). (H) Schematic illustration of the transplantation assay. TA muscle tissues (Host: WT mice) were locally exposed to 18 Gy γ-irradiation and injected with CTX prior to transplantation of either MAS or TA muscle tissue homogenates (Donor: Pax7-YFP mice). Scale bar, 50 μm. Grafted YFP^+^ satellite cells were re-sorted from TA muscles from host mice 14 d following transplantation. qPCR analysis for *Hoxa10* gene expression. (n= 4 mice). Data represent means ± SEM.

Because *Hoxa10* was detected only in limb muscles but not head muscles (Figure 2A), we further determined the expression pattern of *Hoxa10* in muscles throughout the body in adult mice (Figure 2E). qPCR analysis confirmed that all fore- and hind-limb muscles as well as some trunk muscles expressed the *Hoxa10* gene, while it was undetected in head muscles, which is consistent with the *Hoxa10* expression pattern at embryonic developmental regions (Figure 2F). Thus, these results suggest that adult muscles and their satellite cells retain Hox-based positional identities whose distribution almost recapitulates their embryonic origin, which we call “positional memory.”

To test how robust positional memory is maintained in satellite cells, we performed a transplantation study using MAS *(Hoxa10* negative niche) and TA *(Hoxa10* positive niche) muscles from Pax7-YFP knock-in mice *(25)*. Pax7-YFP mouse-derived TA muscle homogenates that contained YFP (Pax7^+^) satellite cells were transplanted into cardiotoxin (CTX)-injected pre-injured MAS muscle (Figure 2G). Immunohistochemical analysis confirmed that TA muscle-derived YFP (Pax7^+^) cells re-populated the MAS niche 2 weeks after transplantation (Figure 2G). Expression of *Hoxa10* mRNA was detected only in MAS muscle tissue transplanted with TA-muscle homogenates (Figure 2G). Next, Pax7-YFP mouse-derived MAS or TA muscle-homogenates were transplanted into TA muscle pre-injured with CTX immediately after 18G γ-ray irradiation of the hindlimbs (Figure 2H). Two weeks following transplantation, immunohistochemistry confirmed that MAS or TA-muscle homogenate-derived YFP (Pax7^+^) cells repopulated the TA muscle niche. Repopulated YFP (Pax7^+^) cells were then re-sorted and qPCR analysis revealed that the higher levels of *Hoxa10* gene expression was detected in re-sorted cells from the TA muscle engrafted with TA-muscle tissues containing YFP (Pax7^+^) cells compared with MAS-muscle tissues containing YFP (Pax7^+^) cells (Figure 2H).

### Postnatal *Hoxa10* deletion in satellite cells decreased region-specific muscle regeneration

Because *Hoxa10* is highly expressed only in limb muscles but not in head muscles, we investigated whether positional memory functions in adult muscle. *Hoxa10*-deficient mice exhibit abnormal morphogenesis of sexual organs, spinal nerves, and femurs during development *(26, 27)*. To investigate the function of *Hoxa10* in satellite cells from adult muscle, we constructed a *Hoxa10*-floxed mouse line (Figure S2A-B). We then generated satellite cell-specific and tamoxifen (TMX)-inducible *Hoxa10* knockout mice (scKO) by crossing *Pax7^CreERT2/+^* mice *(28)* with *Hoxa10*-floxed mice. Genetic inactivation of *Hoxa10* was induced by repeated intraperitoneal injection of TMX in *Pax7^CreERT2/+^;Hoxa10^f/f^* mice. A muscle injury was then induced in both head and hindlimb muscles using a CTX injection (Figure 3A). The number of Pax7^+^ satellite cells were unaltered in TA muscles from scKO mice (Figure 3B). TA muscle mass was significantly reduced in scKO mice compared with those of *Hoxa10^f/f^* (CON) mice; however, MAS muscle was unaffected (Figures 3C–3D), which was consistent with the *Hoxa10* gene expression in limb but not head muscles. CSA of regenerating myofibers in TA but not MAS muscle significantly decreased at both early (5 d, Figure 3E–3F) and late (14 d, Figure 3E and 3G) phases of muscle regeneration following CTX injection.

**Figure 3.**
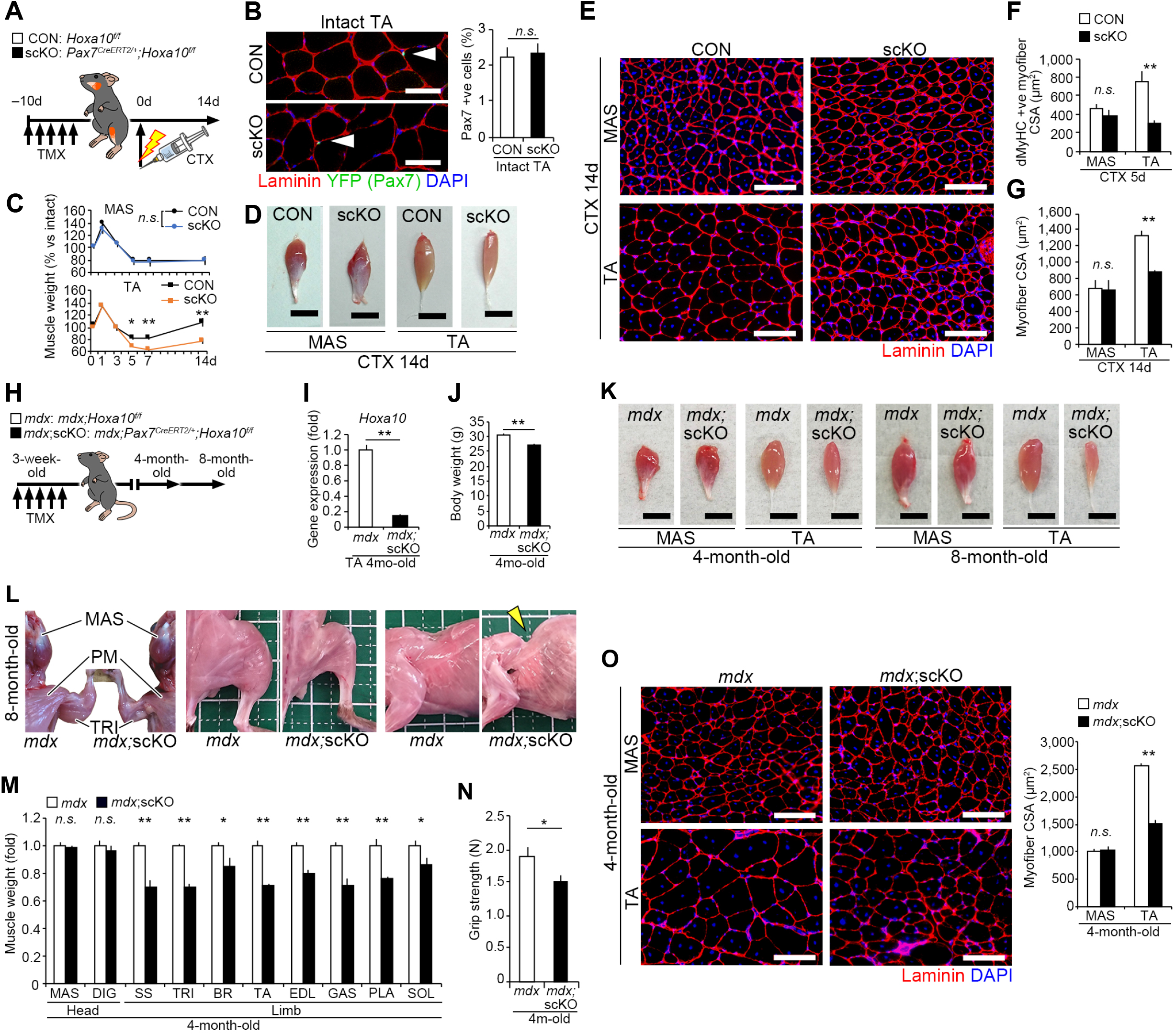
*Hoxa10* inactivation in satellite cells affects muscle regeneration. *Hoxa10* deletion in satellite cells was induced by five-serial intraperitoneal injections of tamoxifen (TMX) in *Pax7^CreERT2/+^;Hoxa10^f/f^* mice before induction of muscle regeneration of MAS and TA muscles by CTX injection. (A) Time-course of TMX and CTX injections. (B) Representative immunohistochemical images of TA muscle tissues for laminin (red), YFP (Pax7, green) and DAPI (blue). Scale bars, 50 μm. Satellite cell number per nucleus (ratio) in muscle cross sections in *Pax7^CreERT2/Pax7-YFP^;Hoxa10^f/f^* (scKO) mice (n= 3 mice each). (C) Weight of regenerated MAS and TA muscles normalized to intact muscles before and after CTX injection (n= 4 mice each). (D) Representative images of regenerated muscles 14 d after CTX injection. Scale bars, 5 mm. (E) Immunohistochemistry for laminin (red) and DAPI (blue) of MAS and TA muscle sections 14 d after CTX injection. CSA of regenerated myofibers were visualized by staining for developmental MyHC (dMyHC) (F) 5 d (TA; n= 5 mice, MAS; n= 4 mice) or for laminin (G) 14 d (n= 5 mice each) after CTX injection. Scale bars, 100 μm. Data represent means ± SEM. (H–O) To perform TMX-inducible ablation of Hoxa10 in satellite cells in *mdx* mice, *mdx;Pax7^CreERT2/+^;Hoxa10^f/f^ (mdx;scKO)* mice were generated. (H) Time-course of TMX injection. TMX treatment was administered at 3 weels of age. (I) qPCR analysis of *Hoxa10* expression in TA muscle (n= 5 mice each). (J) Mean body weight of 4-month-old *mdx* or *mdx;scKO* mice. (K) Representative images of muscles in 4- or 8-month-old mice. Scale bars, 5 mm. (L) At 8 months, *mdx;scKO* mice exhibited kyphosis and severe muscle atrophy throughout the body except in head muscles such as MAS. Grid size (solid line), 1 cm x 1 cm. PM, pectoralis major. (M) Muscle weight of *mdx;scKO* mice normalized to those of *mdx* mice (n= 7 mice each). (N) Grip strength of 4-month-old *mdx* or *mdx;scKO* mice (n= 7 mice each). (O) Immunohistochemistry for laminin (red) and DAPI (blue) to measure myofiber CSA in MAS and TA muscles from 4-month-old mice (n= 5 mice each). Scale bars, 100 μm. Data represent means ± SEM.

We further examined how satellite cell-specific deletion of the *Hoxa10* gene affects muscle regeneration throughout the body in *mdx* mice, which is a Duchenne muscular dystrophy (DMD) model used to monitor chronic muscle degeneration and regeneration throughout the body (Figure 3H). *Hoxa10* inactivation in satellite cells of *mdx* (*mdx*;scKO) mice was induced by a serial injection of TMX at 3 weeks of age and then analyzed at 4 or 8 months of age. *mdx*;scKO mice displayed reduced grip-strength and severe muscle atrophy throughout the body except in head muscles (Figures 3I–3O). At 8 months, the *mdx;scKO* mice showed gradual muscle weakness and noticeable kyphosis, which is a hallmark of extensive and severe muscle wasting in the trunk muscles (Figures 3L). These results indicate that satellite cell-specific deletion of *Hoxa10* perturbed efficient regeneration in limb muscles *in vivo,* demonstrating that a regionally specific gene can control satellite cell function in a site-specific manner.

### *Hoxa10* deletion triggers mitotic catastrophe

Given that satellite cell-specific inactivation of *Hoxa10* caused a severe regeneration defect in limb muscles, even though Hox paralogs are often functionally redundant *(17)*, we tested how satellite cell function is affected by the absence of *Hoxa10 ex vivo.* To inactivate Hoxa10 gene in satellite cells, *Pax7^CreERT2/+^;Hoxa10^f/f^* mice were treated for 5 d with TMX. Individual myofibers from EDL muscle were freshly isolated 5 d after the last injection (designated as 0 h), and then cultured floating in a mitogen-rich media for up to 72 h or plating in growth media (GM) for 6 d (Figure 4A). Immunocytochemistry revealed that the number of Pax7^+^ quiescent satellite cells associated with myofibers was unchanged in the absence of *Hoxa10* (Figure 4B). There was no difference in the total number of Pax7^+^ and/or MyoD^+^ satellite cells per myofiber between CON and scKO at 24 h (Figure 4B), thereby indicating that *Hoxa10* was not required for satellite cell activation. However, after 48 h, the total number of satellite cells per myofiber significantly decreased compared with cells expressing *Hoxa10* (Figure 4B). In adherent culture conditions, we further observed minimal migrating and proliferating satellite cells from myofibers isolated from scKO mice compared with those from CON mice (Figure 4C). These data suggest that *Hoxa10* plays an important role in population expansion of activated satellite cells.

**Figure 4.**
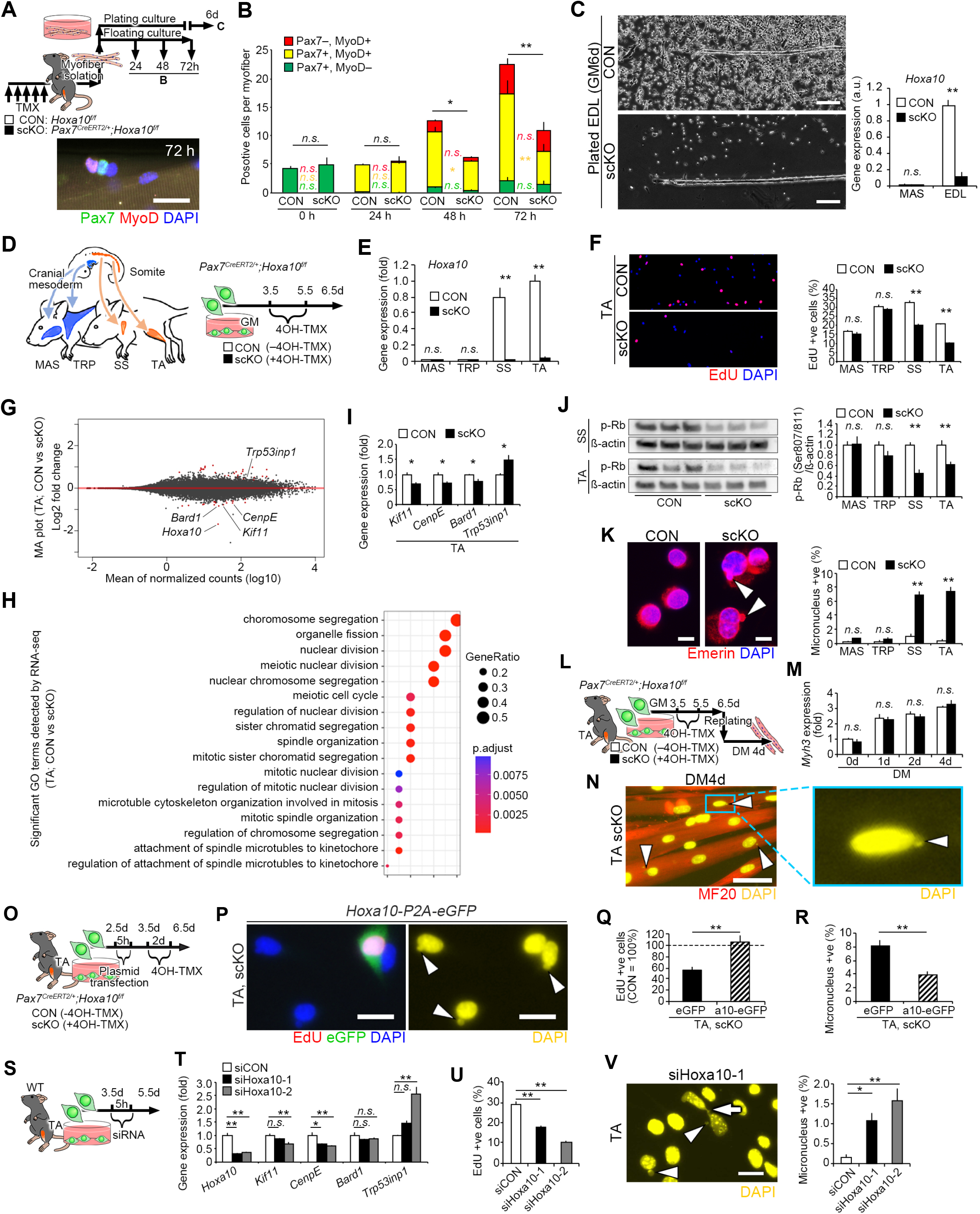
Postnatal inactivation of *Hoxa10* leads to mitotic catastrophe. (A and B) Time schedule of TMX injections used to induce satellite cell specific conditional ablation for Hoxa10 in *Pax7^CreERT2/+^;Hoxa10^f/f^* (scKO) mice. Satellite cells associated with myofibers were isolated from EDL muscle, cultured in floating conditions for up to 72 h, and immunostained for Pax7 and MyoD. (B) The number of Pax7^+^ and/or MyoD^+^ satellite cells per myofiber (n= more than 3 mice, 15–20 myofibers per mouse were counted). (C) Satellite cells associated with myofibers were isolated from EDL and cultured on Matrigel-coated dishes with growth medium (GM) for the adherent condition for up to 6 d. *Hoxa10* gene expression in satellite cells that migrated from the myofibers was determined by qPCR (n= 4 mice each). Data represent means ± SEM. (D-N) To determine the effect of *Hoxa10* in satellite cells, these cells were isolated from the cranial mesoderm-derived (MAS and trapezius (TRP) or somite-derived (SS and TA) muscles of *Pax7^CreERT2/+^;Hoxa10^f/f^* mice and treated with 4OH-tamoxifen (4OH-TMX) in GM culture conditions. (D) Time schedule of 4OH-TMX treatment. (E) qPCR analysis of *Hoxa10* gene expression (n= 6 mice). (F) Proliferation ability of cultured satellite cells determined by 3 h EdU pulse-chase analysis (MAS and TRP, n= 3 mice. SS and TA, n= 6 mice). (G and H) RNA-seq analysis was performed in satellite cells isolated from TA muscle. Mean MA plots depicting fold-change in gene expression (G) and significant Gene Ontology (GO) terms (H) (n= 9 mice). (I) qPCR analysis of mitosis and chromosome segregation-related gene expression (n= 6 mice). (J) Immunoblot analysis for the expression of phosphorylated (Ser807/811) Rb in satellite cells from scKO mice normalized phosphorylated levels from control mice (n= 6 mice). (K) Micronuclei were visualized by staining for Emerin (red) and DAPI (blue) in satellite cells isolated from EDL (e.g. limb muscle). Arrowheads show micronuclei. Scale bar, 20 μm. Micronucleus^+^ cells were quantified (n= 5 mice). Data represent means ± SEM. (L-N) To examine the effect of *Hoxa10* on myogenesis, satellite cells were isolated from TA muscle of *Pax7^CreERT2/+^;Hoxa10^f/f^* mice, treated with 4OH-TMX, and induced to differentiate with differentiation medium (DM) for 4 d. (L) Time-course for 4OH-TMX treatment and differentiation. (M) qPCR analysis of Myh3 gene expression (n= 3 mice). (N) Representative immunostaining of myotubes using MF20 (MyHC) and DAPI. Arrowheads show micronuclei in myotubes. Scale bar, 50 μm. Data represent means ± SEM. (O-R) To determine the effect of exogenous expression of Hoxa10 in satellite cells, satellite cells were isolated from TA or MAS muscles of *Pax7^CreERT2/+^;Hoxa10^f/f^* mice, transfected with *pCMV-Hoxa10-P2a-eGFP* or *pCMV-eGFP* (control) vectors, treated with 4OH-TMX, and analyzed 6.5 d after isolation. (O) Time schedule of vector transfection and 4OH-TMX treatment. (P) Representative images for eGFP (green), EdU (red), and DAPI (blue) staining in TA-derived scKO cells transfected with *pCMV-Hoxa10-P2A-eGFP* vector. Arrowheads show micronuclei in untransfected (eGFP^−^) cells. Scale bar, 20 μm. EdU^+^ cells (Q) or micronucleus^+^ cells (R) were quantified (n= 4 mice). Data represent means ± SEM. (S–V) To investigate the effect of reduced *Hoxa10* levels on satellite cells; satellite cells were isolated from TA muscles of WT mice and transfected with siRNA against *Hoxa10.* (S) Experimental timeline for siRNA transfection. (T) Gene expression profiles were determined with qPCR (n= 4 mice). (U) Quantification of EdU^+^ cells in DAPI^+^ cells (n= 4 mice). (V) A representative fluorescence image of satellite cells stained with DAPI (yellow). Arrow, chromosome bridge; arrowheads, micronucleus. Scale bar, 20 μm. Micronucleus^+^ cells were quantified (n= 4 mice). Data represent means ± SEM.

To further evaluate the effect of *Hoxa10* inactivation on satellite cell function, we used the developmentally different muscles, such as TRP and MAS muscles that originate from the cranial mesoderm and SS and TA muscles with a somite origin (Figures 4D and S3A). These cell types are clearly distinguishable by their *Hoxa10* expression levels (Figure 4E). A cell cycle assay including EdU pulse-chase analysis revealed that proliferation was impaired in both SS and TA-derived scKO satellite cells, while both TRP and MAS-derived scKO satellite cells were unaffected (Figures 4F and S3B-S3D, and S3F-S3G). A terminal deoxynucleotidyl transferase dUTP nick end labeling (TUNEL) assay showed that apoptosis was not involved in the cell-number decline observed in scKO TA satellite cells (Figure S3H).

Given that proliferation in *Hoxa10*-deficient satellite cells was decreased compared with those expressing *Hoxa10,* we next examined how population expansion of limb-derived satellite cells was impaired by loss of Hoxa10. RNA-seq analysis revealed that mitosis-related gene sets for chromosomal segregation, meiotic nuclear division, and spindle organization significantly changed in *Hoxa10* deficient satellite cells compared with CON cells (Figures 4G and 4H). *Kif11, CenpE,* and *Bard1* regulate mitotic spindle formation and genomic stability, and *Trp53inp1* is upregulated by cellular stress and induces cell-cycle arrest. qPCR analysis also indicated significant changes in the expression of these genes in TA-derived scKO satellite cells (Figure 4I). Dephosphorylation of Rb, which is known to be induced by genomic instability, arrests the cell-cycle. Our immunoblot analysis confirmed that *Hoxa10* ablation led to a significant decrease in the level of phosphorylated Rb in limb-derived scKO satellite cells (Figure 4J). Micronuclei and chromosomal bridges are hallmarks of chromosome misalignment and mitotic catastrophe. We found that the ratio of micronuclei and chromosomal bridges increased in only somite-derived scKO satellite cells and not cranial mesoderm-derived scKO satellite cells (Figures 4K, S3E, and S3I-S3J). It should be noted that genetic ablation of *Hoxa10* did not influence myogenic differentiation or selfrenewal of TA-derived satellite cells, but that differentiating myotubes contained abundant micronuclei (Figures 4L–4N, and S3K-S3N). Importantly, the decreased proportion of EdU^+^ proliferative cells and the increased proportion of micronucleus^+^ cells in somite-derived scKO cells were rescued by *pCMV-Hoxa10-P2A-eGFP* vector-mediated exogenous Hoxa10 expression in TA-derived satellite cells (Figures 4O–4R). These findings in our genetic ablation models were confirmed by siRNA-mediated knockdown of Hoxa10 in limb-derived satellite cells in WT mice (Figures 4S–4V and S3P-S3R). Taken together, our data indicate that postnatal reduction of *Hoxa10* in somite-derived satellite cells led to genomic instability and mitotic catastrophe, resulting in a reduction in proliferation ability, probably through abnormal expression of genes that control chromosome segregation and mitosis.

### *HOXA10* expression and function are conserved in human satellite cells

We evaluated if the function of *HOXA10* characterized in mice was conserved in satellite cells from adult humans. We used semitendinosus (ST) muscle from lower limb and MAS and buccinator (BUC) muscles from the head (Figure 5A) of adult humans. qPCR analysis confirmed that *PAX3* or *TCF21* genes were detected in ST limb or MAS head muscles, respectively (Figure 5B), similar to the results from mouse satellite cells *(6)*. We showed that the *HOXA10* gene was detectable in ST muscle and in its associated satellite cells (Figures 5B–5C). It is likely that *HOXA10* gene expression is robustly maintained in ST-satellite cells because long-term passaging did not influence its expression level (Figure 5D).

**Figure 5.**
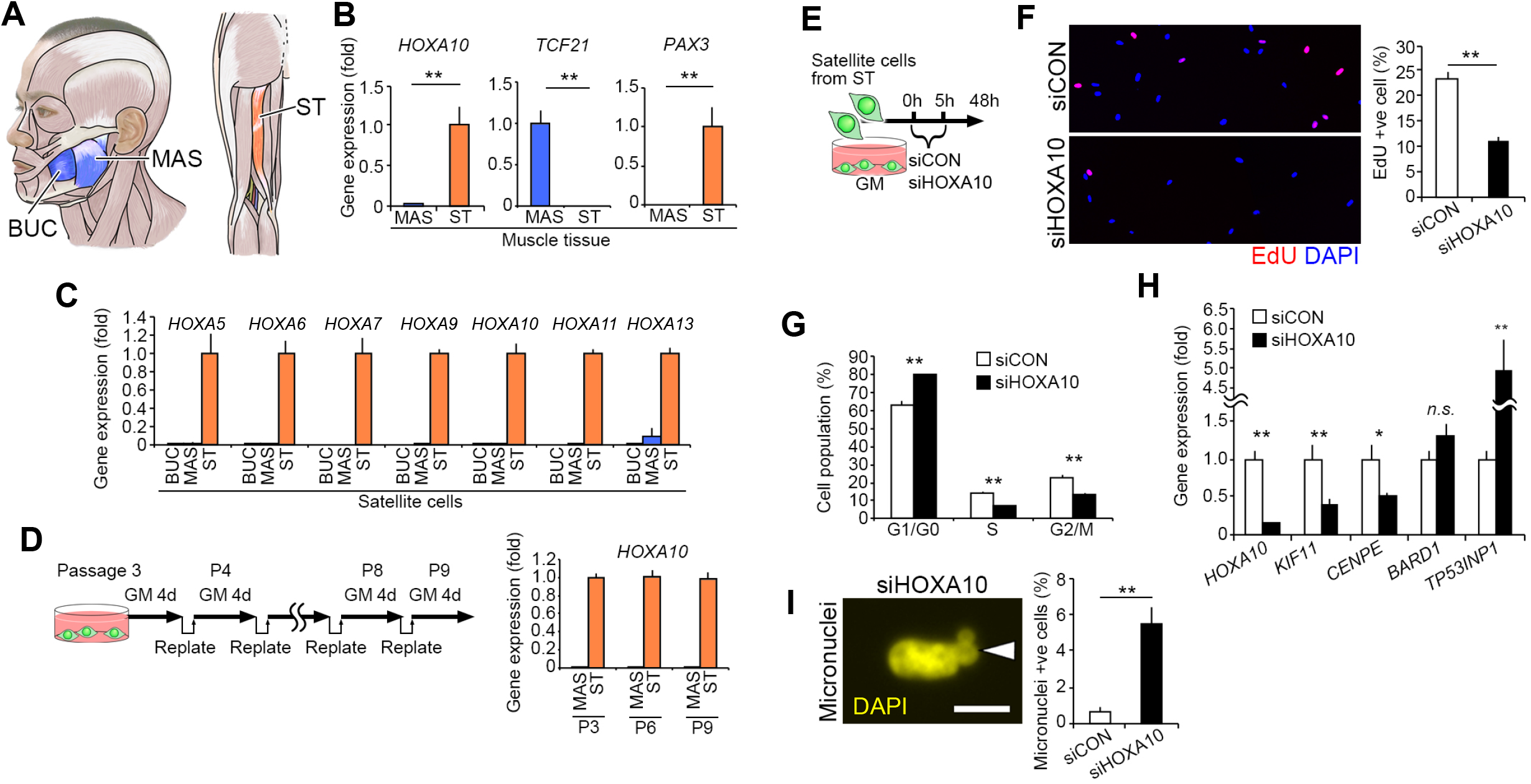
Expression and function of *HOXA10* in human satellite cells. (A-D) Satellite cells were isolated from the lower limb semitendinosus (ST) muscle, and MAS and buccinator (BUC) muscles from heads of adult humans (A). (B and C) qPCR analysis of gene expression in skeletal muscle tissues (B) (n= 4 individuals for each muscle) and satellite cells (C) (BUC, MAS, and ST: n= 3, n= 4, and n= 3 individuals, respectively). Data represent means ± SEM. (D) Satellite cells isolated from MAS and ST muscles were repeatedly passaged for up to 9 passages in culture. Samples were collected at the 3^rd^, 6^th^ and 9^th^ passages and HOXA10 gene expression was determined by qPCR. Data represent means ± SEM. (E–I) To examine the effect of *HOXA10* knockdown, siRNA targeting *HOXA10* was transfected into satellite cells derived from ST muscle *in vitro.* (E) Time schedule of siRNA transfection. (F) Quantification of EdU^+^ cells (n= 4 individuals). (G) Distribution of cells in different cell-cycle phases were evaluated based on DNA content (n= 3 individuals). (H) qPCR analysis of mitosis and chromosome segregation-related gene expression (n= 12 individuals). (I) A representative image of micronucleus^+^ satellite cells (n= 4 individuals). Data represent means ± SEM.

To further examine the function of *HOXA10* in ST-derived satellite cells, siRNA-mediated knockdown of HOXA10 was performed (Figure 5E). Consistent with the mouse phenotypes, reducing *HOXA10* expression levels resulted in the impairment of proliferation ability, as well as a downregulation in the expression of *KIF11* and *CENPE* gene in limb satellite cells, while *TP53INP1* was upregulated compared with the control siRNA-transfected cells (Figure 5F–5H). Notably, we confirmed that the ratio of micronuclei increased in HOXA10 knocked-down satellite cells (Figures 5I). Thus, we conclude that *HOXA10* is expressed in limb satellite cells and functions in population expansion of satellite cells in human limb muscle.

## DISCUSSION

Expression of region-specific genes including Hox genes has been observed in muscles both during embryonic development and in adult mice *(6, 9, 15, 21–23, 29, 30)*. However, whether regional specific genes have a function in satellite cells of adult muscle remains to be elucidated. In the present study, our comprehensive analysis demonstrated that Hox gene expression profiles in adult muscles almost mirror embryonic origin. These regionally specific Hox expression patterns were regulated epigenetically, which could act as a molecular signature that reflects the embryonic history in both adult skeletal muscles and resident muscle cells. Moreover, our results demonstrated that Hox gene expression displayed functional memory that governed stem cell function in adult muscle in mice and humans.

A recent study reported that DNA hypermethylation on the HOX-A locus is positively correlated with upregulation of HOX-A cluster genes in human glioblastoma *(31)*. This correlation suggests that DNA hypermethylation permits upregulation of HOX-A cluster genes. Consistent with this finding, our DNA methylome analysis revealed hypermethylation of the Hox-A locus in adult limb muscles and its associated satellite cells. Thus, we speculated that expression of the limb-specific Hox-A cluster genes was predominantly regulated through DNA hypermethylation. In addition to DNA methylation, histone modifications may be also involved in regulating Hox-A cluster genes in satellite cells *(32)*. We observed that the expression levels of Hox-A to −D cluster genes in limb-derived satellite cells were almost similar between young and aged mice (data not shown), even though DNA methylation status in satellite cells alters globally with aging *(33)*. In contrast to our results, a recent study described the upregulation of *Hoxa9* in limb satellite cells during aging, and upregulated *Hoxa9* perturbed cell-cycle entry through the misregulation of genes involved in development, such as Wnt, TGF-β, and JAK/STAT pathways *(23)*. Because our data demonstrated that genetic loss of *Hoxa10* in satellite cells of young adult mice resulted in a decrease in proliferation with genomic instability, *Hoxa9* and *Hoxa10* have entirely distinct function in satellite cells even though both are adjacent in the Hox-A cluster genes.

Muscular dystrophies are characterized by progressive muscle weakness and wasting, although they vary in severity and body-regions affected *(8)*. Patients with DMD are preferentially inflicted with a severe pathology in limb and trunk muscles, but not in craniofacial muscles, particularly eye muscles. Emery-Dreifuss muscular dystrophy (EDMD) patients show progressive muscle weakness, especially in the arms and lower legs. Moreover, particular muscles are known to be affected in oculopharyngeal muscular dystrophy (OPMD), limb girdle muscular dystrophy (LGMD), and facioscapulohumeral muscular dystrophy (FSHD) patients. This body-region specific muscle pathology is also observed during aging and microgravity. For instance, MAS muscle from the head is resistant to atrophy during aging and spaceflight *(34, 35)*, whereas we observed that an acute muscle injury promoted a poor regenerative ability in the head muscle compared with the efficient regeneration of the limb muscle *(36)*. In this study, we identified that *Hoxa10* provided positional memory of limb muscles, thereby regulating genomic stability and expansion of satellite cell populations. Based on these data, functional positional memory may explain the varying regenerative capacity between different muscle groups and account for specific regions of muscles to be susceptible to diseases. Further studies will elucidate the link between positional memory and region-specific pathologies in muscle disorders.

In conclusion, we demonstrated that muscles and their associated satellite cells retain their own positional memory based on developmental origin and anatomical location, which dictates stem cell function in a regionally specific manner in adults. Although our results suggest that niche-exposure at least for 2 weeks following ectopic transplantation did not influence Hox-based positional memory in satellite cells, the effect of a longer period exposure to different niche remains unclear. Furthermore, when and how the positional memory is established during embryogenesis and robustly maintained in adult tissues remain to be determined. Our findings indicate that Hox-based functional memory in satellite cells potentially regulates topographical gene expression in adult skeletal muscles, providing a molecular basis for a body region-specific pathogenesis in muscle weakness related to age-related sarcopenia as well as muscular dystrophies.

## MATERIALS AND METHODS

### Animals

The Experimental Animal Care and Use Committee of Nagasaki University and Kumamoto University approved animal experimentation (Ref. No. 1203190970 and A30-098). The BRUCE-4 ES cell line (C57/BL6J) was used to generate the *Hoxa*10-floxed mouse line. A targeting vector was generated to modify the *Hoxa10* locus by inserting *loxP* sequences upstream of exon 2 and downstream of exon 3 in the *Hoxa10* gene (Figure S2A). To induce conditional ablation of the *Hoxa10* gene, *Hoxa10*-floxed mice were crossed with *Pax7^CreERT2/+^ (28)* mice. *Pax7-YFP* (*Pax7^YFP/YFP^*) mice were used to visualize satellite cells expressing Pax7-YFP fusion proteins *(25)*. *mdx* mice were obtained from CREA Japan, Inc (Tokyo, Japan) and were crossed with scKO mice to generate *mdx*;scKO mice. In *Pax7^CreERT2/+^;Hoxa10^f/f^* mice, inactivation of *Hoxa10* was induced by repeated intraperitoneal injection of tamoxifen (TMX) dissolved in corn oil (5 μL/g, 20 mg/mL) as previously described *(37)*.

For muscle injury, mice were anesthetized and the hair on the lower jaw and lower leg areas was removed with a depilatory cream (Epilat, Kracie). The facial nerve was percutaneously identified to avoid needle damage. To induce muscle injury, 10 μM cardiotoxin (CTX, Sigma-Aldrich) (50 μL for TA and 40 μL for MAS) was injected into the muscles of anesthetized mice. CTX-injected and non-injected (intact) muscles were harvested for transverse sectioning and immunostaining. For cell transplantation, mice were anesthetized and 18 Gy of γ-radiation were exposed to the hind legs to prevent population expansion of endogenous satellite cells. The rest of the body was protected by a 2-cm-thick lead plate.

### Methylome analysis

For Microarray-based Integrated Analysis of Methylation by Isoschizomers (MIAMI) analysis, a genome-wide analysis of DNA methylation was performed based on the protocol (http://epigenome.dept.showa.gunmau.ac.jp/~hatada/miami/image/MIAMI%20Protocol%20V4.pdf *(18)*) as descried previously *(38)*. Briefly, genomic DNA was extracted from MAS and TA muscle tissues of mice via the standard proteinase K method. Purified genomic DNA was digested with methylation-sensitive *HpaII* and methylation-insensitive *MspI*, followed by adaptor ligation and PCR amplification. Amplified DNA from MAS and TA tissues was labeled with Cy3 and Cy5, respectively. Labeled DNA was then co-hybridized using the Oligo aCGHZChIP-on-chip Hybridization kit (Agilent Technologies, CA) and the difference in the *HpaIIZMspI* signal ratio was determined as the methylation value. Values of <0.5 and >2 denoted DNA hypomethylation and hypermethylation, respectively.

For further methylome analysis, the Post-Bisulfite Adaptor Tagging method (PBAT)-mediated whole-genome bisulfite sequencing was performed as previously described *(19, 20)*. For preparation of PBAT libraries, genomic DNAs were extracted from MAS and TA muscle tissues and plated satellite cells of adult mice. Sequencing was performed by Macrogen Japan Corp. (Kyoto, Japan) using the HiSeq X Ten, and the reads were mapped to the mouse mm10 reference genome using BMap (http://itolab.med.kyushu-u.ac.jp/BMap/index.html) with default parameter settings. Methylation levels of CG sites were calculated for only those covered by five or more reads. Methylation levels of 1-kb sliding sliding windows, promoters (<2 kb upstream of TSS), gene bodies, CGIs, and TSS-proximal regions were determined by averaging the methylation levels of CG sites in individual features and were analyzed using two unsupervised classification methods, namely hierarchical clustering and principal-component analysis (PCA). DMRs were identified using the program metilene *(39)* with default parameters. DMRs were filtered so that they have at least 5 CpGs with a q-value less than 0.05 and methylation difference larger than 20%. The nearest neighbor gene of a DMR was defined as the gene associated with the DMR. All functional enrichment analysis with gene ontology (GO) biological process terms was performed using TopGO (R package ver. 2). A p-value of 0.01 was set as the significance level for gene set enrichment with an additional requirement that at least 5 genes are involved in the annotated biological function. Raw DNA methylation data sets are available from the GEO public depository under the GSO accession number (GSE154056).

### Statistical analysis

Statistical analysis was performed in Microsoft Excel or Prism 8 (GraphPad Software, La Jolla, CA). For statistical comparisons of two conditions, the Student’s t-test was used. For comparisons of more than two groups, the data were analyzed with one-way or two-way ANOVA, according to the experimental design, followed by Sidak’s multiple comparison test. *P*-values of <0.05 (*), <0.01 (**) were considered statistically significant. Results are presented as means ± SEM. *n.s.* indicates results that were not statistically significant.

## SUPPLEMENTARY MATERIALS

Supplementary materials include supplementary materials and methods, three figures, and one table.

## ACKNOWLEDGMENTS

We thank all lab members for their technical support. We thank Deneen Wellik, Peter Zammit, Shin’ichi Takeda, and Shahragim Tajbakhsh for fruitful discussions and helpful comments. This work was supported by the Japan Agency for Medical Research and Development (AMED, 16bm0704010h0001; 18ek0109383h0001, and 19bm0704036h0001), and the Grant-in-Aid for Scientific Research JSPS KAKENHI (18H03193, 18K19749, and 16J09948). This work was also supported, in part, by the Takeda Science Foundation, Japanese Physical Therapy Association, the International Research Core for Stem Cell-based Developmental Medicine (Kumamoto University), the Cooperative Research Project Program of the Medical Institute of Bioregulation (Kyushu University), and the Platform Project for Supporting Drug Discovery and Life Science Research (Basis for Supporting Innovative Drug Discovery and Life Science Research (BINDS)) from AMED under Grant Number JP20am0101103 (support number 1963).

## AUTHOR CONTRIBUTIONS

K.Y. designed and performed the experiments, analyzed the data, and wrote the manuscript. H.N., Y.Ki., D.S., J.N., Y.T., Y.Ka., F.M. H.A., and Y.Oh performed the experiments and analyzed the data. N.O., A.Y., S.O., Y.S., T.S., K.C., K.I., I.A., Y.Og and T.I. provided analytical tools and gave technical support. Y.On conceived the project, designed and performed the experiments, analyzed the data, assembled the input data, and wrote the main manuscript. All authors read and approved the final manuscript.

